# A comparative network analysis of high-risk MYCN amplified and high-risk MYCN non-amplified neuroblastoma patients

**DOI:** 10.1101/2024.10.24.620136

**Authors:** Erin Chang, Charissa Chou, Wen-I Chang, Joshua N Honeyman, Alper Uzun

## Abstract

Neuroblastoma is the most common extracranial solid tumor in children. Under contemporary staging guidelines, amplification of MYCN, defined as possessing 10 or more copies of the MYCN gene, is sufficient to classify a patient as high-risk. We aim to investigate the complex genetic landscape of neuroblastoma through a network biology perspective, focusing on complex protein-protein interaction (PPI) networks. Reanalyzing three historical neuroblastoma RNA-Seq datasets, we conducted a comparative network analysis of high-risk MYCN amplified patients and high-risk MYCN non-amplified patients to identify PPI networks that serve as predictive biomarkers for developing high-risk disease. We used R scripts to extract the top 100 most highly expressed genes from each dataset and then analyzed the expression profiles using Proteinarium, a network analysis tool that identifies clusters of patients with shared PPI networks. Statistically significant clusters were annotated and analyzed for network similarity between datasets. Our study isolated four significant clusters with majority high-risk MYCN amplified patients. We annotated the genes unique to high-risk MYCN amplified patients and identified several potential network biomarkers for this particular genotype of neuroblastoma, including ribosome biogenesis (RPL7A, RPL30, RPL35, RPS8), heat shock protein (HSPA8, HSP90AA1, HSP90AB1), matrix metallopeptidase (MMP2, MMP9, MMP13), and collagen (COL2A1, COL5A3, COL6A2, COL11A1, COL16A1) gene networks. The results of this study provide potential therapeutic targets for high-risk MYCN amplified neuroblastoma.

## Introduction

Neuroblastoma (NBL) is the most common extra-cranial solid tumor of childhood. Despite progress in low-risk and intermediate-risk populations, high-risk patients retain a poor 5-year survival rate of 40-50% (1), indicating an urgent need for therapies targeting high-risk disease mechanisms. Amplification of the *MYCN* gene is a strong prognostic marker for high-risk neuroblastoma according to the International Neuroblastoma Risk Group Staging System (INRGSS) staging guidelines (2). Thus, we elected to identify biomarkers for the specific subgroup of high-risk *MYCN* amplified neuroblastoma patients.

Previous studies report a low mutation burden among high-risk neuroblastoma patients with a median exonic mutation frequency of 0.60 per megabase (3). However, like all other cancers, neuroblastoma is a complex disease resulting from the cumulative effect of various interacting genetic mutations. Existing studies that search for prognostic markers among these mutations largely focus on the differential expression of single genes, limiting the scope of potential targeted therapeutics. Given the complex genetic landscape of high-risk neuroblastoma, we used a network biology approach, identifying complex protein-protein interaction (PPI) networks rather than individual proteins as biomarkers for poor prognosis in high-risk *MYCN* amplified neuroblastoma (4).

In this study, we conducted a comparative network analysis of high-risk *MYCN* amplified and high-risk *MYCN* non-amplified patients, utilizing the novel network analysis tool Proteinarium to identify PPI networks that serve as predictive biomarkers for poor prognosis in high-risk *MYCN* amplified neuroblastoma. We analyzed gene expression data for a high-risk *MYCN* amplified cohort and high-risk non-amplified cohort and isolated the PPI networks exclusive to the amplified cohort. These PPI networks are potentially therapeutically targetable.

## Materials and Methods

We used three datasets for our analysis. The NCI Therapeutically Applicable Research to Generate Effective Treatments (TARGET) dataset contained whole exome sequencing data from 1,076 patients (dbGaP sub-study ID: phs000467). The Sequencing Quality Control Consortium dataset (GSE49711) contains RNAseq gene expression data from 498 patients (5). These datasets were obtained from two repositories (respectively): cBioPortal and the Gene Expression Omnibus (6-8). In addition to the two patient-derived datasets, we also utilized gene expression data derived from 41 neuroblastoma cell lines that was also made available through the Gene Expression Omnibus (GSE89413) (9). The TARGET and GSE49711 datasets provided patient clinical data including risk category and death from disease, which we used for a survival analysis. The TARGET dataset included overall survival in months, allowing us to create Kaplan-Meier survival plots for the patients analyzed from that dataset.

For the TARGET and GSE49711 datasets, we specifically focused on patient samples classified as high-risk due to the distinct genetic characteristics and worse survival outcomes of this group. In the TARGET dataset, which includes 1,076 patients, 249 had complete RNA sequencing data. From this group, there were 213 high-risk patients, 68 with *MYCN* amplification and 145 with non-amplified *MYCN*. In the GSE49711 dataset, 498 patients had full RNA sequencing data. There was a total of 175 high-risk patients. Of these, 92 were *MYCN* amplified and 83 were *MYCN* non-amplified. The GSE89413 dataset included RNA sequencing data on 27 *MYCN* amplified and 14 *MYCN* non-amplified neuroblastoma cell lines. The inclusion of these cell-line data allowed us to compare human samples and gain insights into which PPI networks could potentially be targeted in in-vitro experiments.

We used R version 4.1.0 in RStudio to create custom scripts that filtered out noncoding genes (such as LINC genes and open reading frame genes) and extracted the top 100 most highly expressed genes from both high-risk *MYCN* amplified and high-risk *MYCN* non-amplified patients in each dataset. These gene lists, representing high-risk *MYCN* amplified and non-amplified samples, were then uploaded to Proteinarium for network analysis. Proteinarium converts user defined seed genes to protein symbols and maps them onto the STRING interactome provided by version 11 of the STRING database (10). The program then builds PPI networks for each sample and uses the Fisher-Exact test to identify significant clusters of patients with shared PPI networks (11).

For our analysis in Proteinarium, we used the default settings, which included:

- Max Number of Vertices to Render: 50
- Meta Cluster Threshold: 0.8
- Number of Bootstrapping Rounds: 0
- Max Path Length: 3
- Repulsion Constant: 1.2

We ran the high-risk *MYCN* amplified patients as Group 1 and the high-risk *MYCN* non-amplified patients as Group 2. The Proteinarium workflow is depicted in Figure 1.

**Figure 1.**
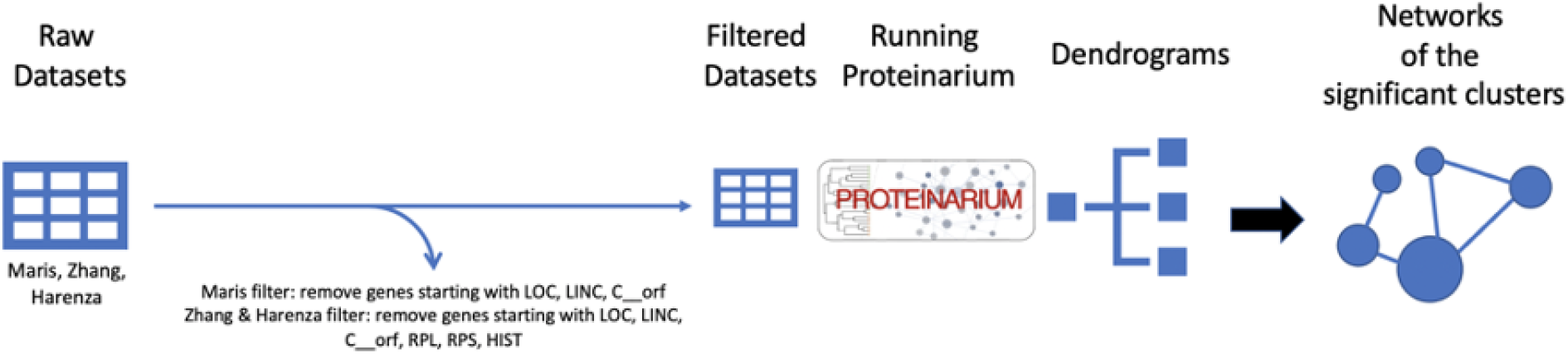
Workflow diagram outlining our network analysis process, including filtering raw datasets, running datasets through Proteinarium, and obtaining significant clusters.

After identifying the four clusters of interest, we focused on the gene sets specific to the *MYCN* amplified patients in these clusters. To annotate the networks from our chosen clusters, we uploaded the *MYCN* amplified gene sets to g:Profiler. g:Profiler’s functional annotation tool enables users to consolidate an uploaded gene list’s most relevant gene ontology terms, including biological processes, cellular components, molecular functions, and KEGG pathways (12). We used the false discovery rate (FDR) with a significance threshold of 0.01 to determine significance of listed processes and pathways from g:Profiler.

Lastly, we used R scripts to conduct separation tests of each cluster against itself and the others in order to measure the average distances between two networks in the STRING interactome, a set of protein-protein interactions composed of 141,296 experimentally determined physical interactions between 13,460 proteins (13). A positive separation test score indicates topological separation between two networks in the interactome, suggesting that the two networks are biologically distinct. A negative score indicates topological overlap in the interaction, suggesting that the two networks are biologically similar. There is no standard value indicating complete network overlap, but running a network against itself gives us a negative value that is useful as a relative comparison to show what a complete overlap looks like for that specific network. We ran ten separation tests in total.

## Results

### Protein-Protein Interaction Network Mapping with Proteinarium

Each of the three datasets produced a unique dendrogram, a layered graph that organizes user-uploaded samples into clusters according to their PPI network similarity. Figures 2-4 depict the dendrograms generated from each dataset, visualized using Interactive Tree of Life (iTOL) software (14). *MYCN* amplified and *MYCN* non-amplified samples tended to cluster together in all three dendrograms. We selected four clusters composed of majority *MYCN* amplified samples for network analysis, as indicated in the highlighted sections of Figures 2-4.

**Figure 2.**
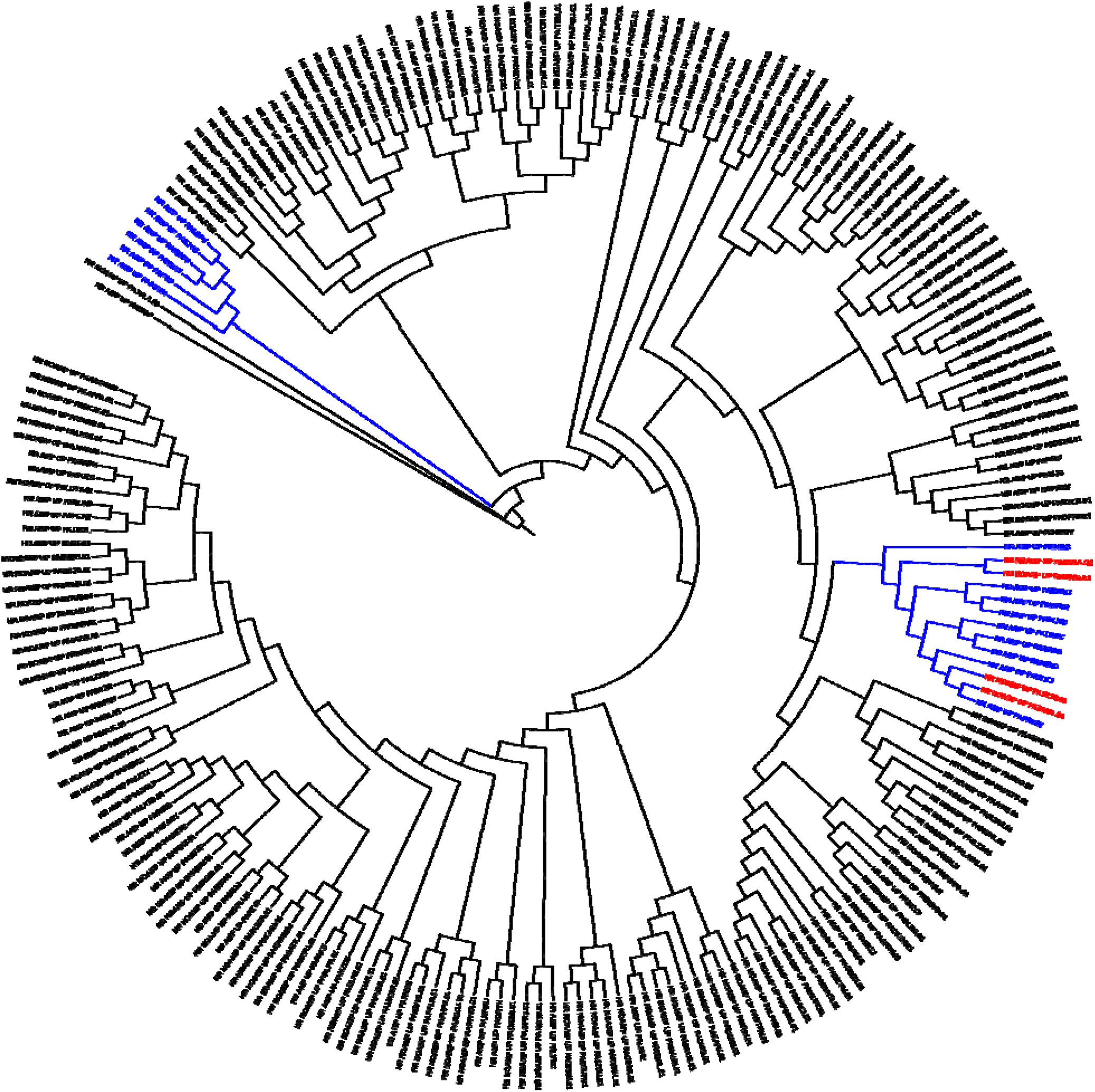
Circular visualization of the TARGET dataset’s dendrogram. Cluster A is highlighted in blue on the left side of the figure. Cluster B is highlighted in blue and red on the right side of the figure. Blue leaves of the dendrogram indicate a *MYCN* amplified sample, while red indicates a *MYCN* non-amplified sample. Cluster A demonstrates a significant shared PPI network among 6 *MYCN* amplified patient samples. Cluster B contains a significant shared PPI network among 9 *MYCN* amplified patient samples and 3 *MYCN* non-amplified samples.

**Figure 3.**
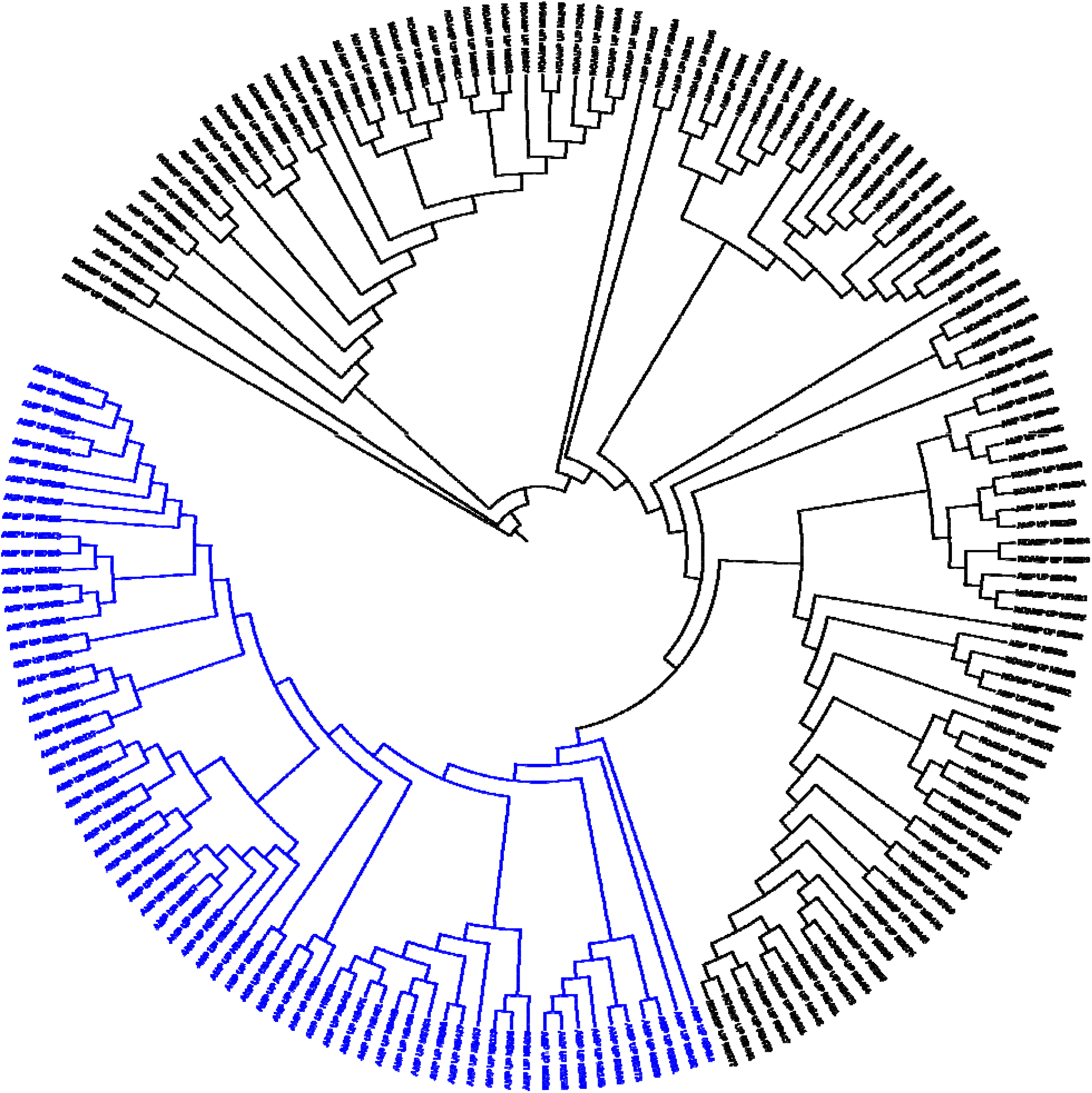
Circular visualization of the GSE49711 dataset’s dendrogram. Cluster C is highlighted in blue. This cluster contains a significant shared PPI network among 65 *MYCN* amplified patient samples.

**Figure 4.**
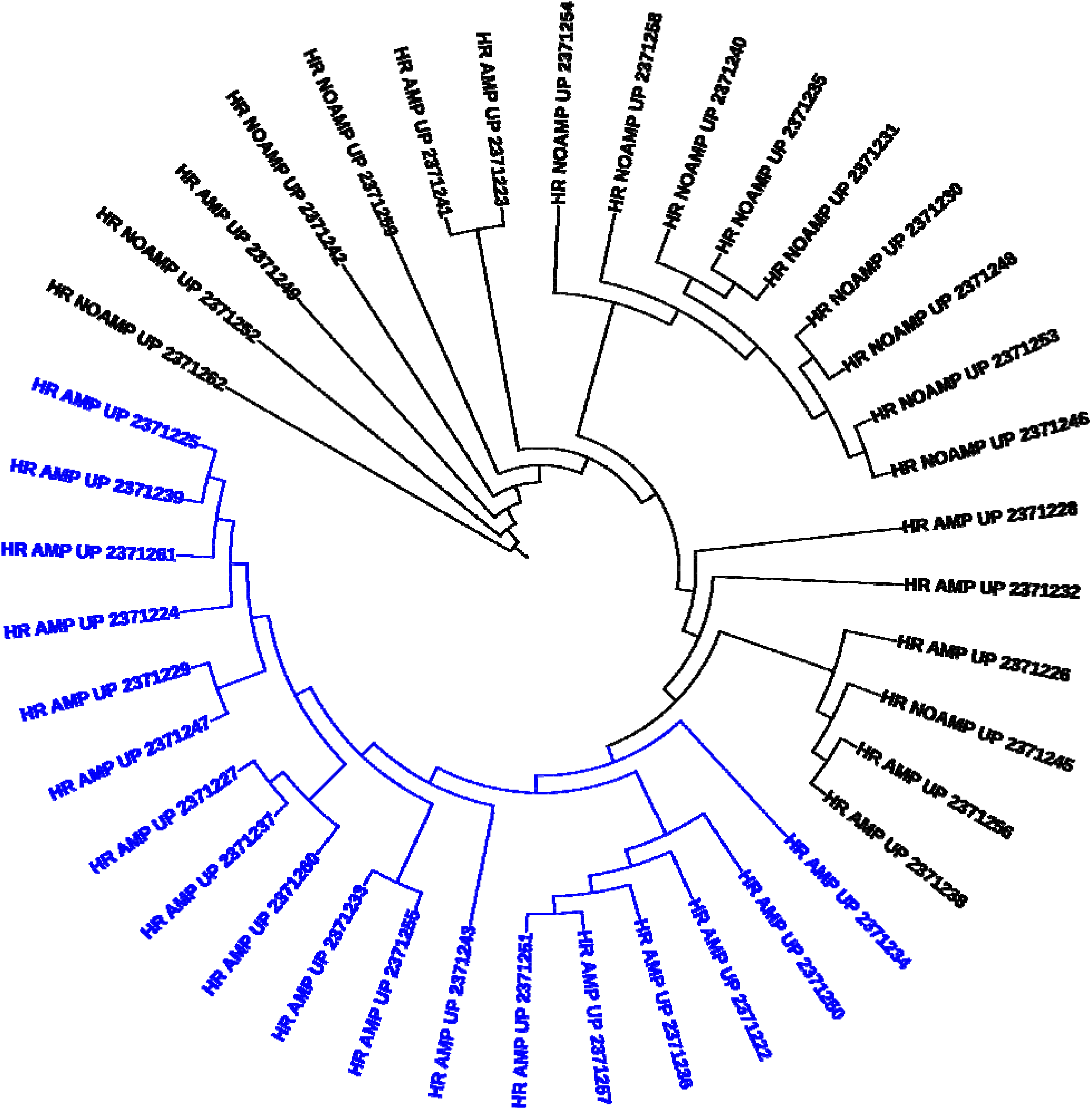
Circular visualization of the GSE89413 dataset’s dendrogram. Cluster D is highlighted in blue. This cluster contains a significant shared PPI network among 18 *MYCN* amplified cell-line samples

Proteinarium uses the Fisher-Exact test to determine which patient clusters have a significant degree of network similarity. Using a significance threshold of 0.05, we isolated four clusters with majority high-risk *MYCN* amplified samples for further analysis. These clusters were C208 and C121 from the TARGET dataset (Clusters A and B, respectively), C11 from the GSE49711 dataset (Cluster C), and C11 from the GSE89413 dataset (Cluster D). Table 1 provides a description of each cluster, including the dataset used, the genes removed when running Proteinarium, the p-value of the cluster, and the number of *MYCN* amplified patients and *MYCN* non-amplified patients.

**Table 1.** Summary of each cluster identified in our network analysis, including the dataset used, the genes removed when running Proteinarium, the p-value of the cluster, and the number of *MYCN* amplified patients and *MYCN* non-amplified patients.

| Cluster | Dataset | Genes Removed | p-value | MYCN amplified samples | MYCN non-amplified samples |
| --- | --- | --- | --- | --- | --- |
| A | TARGET | Noncoding genes | 0.000906 | 6 | 0 |
| B | TARGET | Noncoding genes, RPL/RPS genes, HIST genes | 0.002093 | 9 | 4 |
| C | GSE49711 | Noncoding genes, RPL/RPS genes, HIST genes | < 0.0001 | 65 | 0 |
| D | GSE89413 | Noncoding genes, RPL/RPS genes, HIST genes | 0.000022 | 18 | 0 |

We identified several potential network biomarkers for high-risk *MYCN* amplified neuroblastoma, including ribosome biogenesis (*RPL7A, RPL30, RPL35, RPS8*), collagen (*COL11A1, COL16A1, COL2A1, COL5A3, COL6A2*), matrix metallopeptidase (*MMP2, MMP9, MMP13*), and heat shock protein (*HSPA8, HSP90AA1, HSP90AB1*) networks. These PPI networks are depicted in Figures 5-8, which were produced with Gephi. We applied Gephi’s modularity tool, which organizes nodes into colored modules with a high density of connections and customized the node sizes to reflect their relative number of connections to other nodes. Figure 5 illustrates the network of Cluster A, which contains a network of various ribosome biogenesis hub genes within the green module. In Figure 6, the Cluster B network exhibits a network composed of several collagen hub genes within the green and blue modules. In Figure 7, several hub genes including *GNB2L1, GRB2, HSPA2, HSP90AA1, UBA52, UBB, ACTB*, and *DYNLL2* dominate Cluster C. Many of the same hub genes appear in Cluster D, including *GNB2L1, GRB2, HSPA2, HSP90AA1, UBA52, UBB*, and *ACTB*, as indicated in Figure 8.

**Figure 5.**
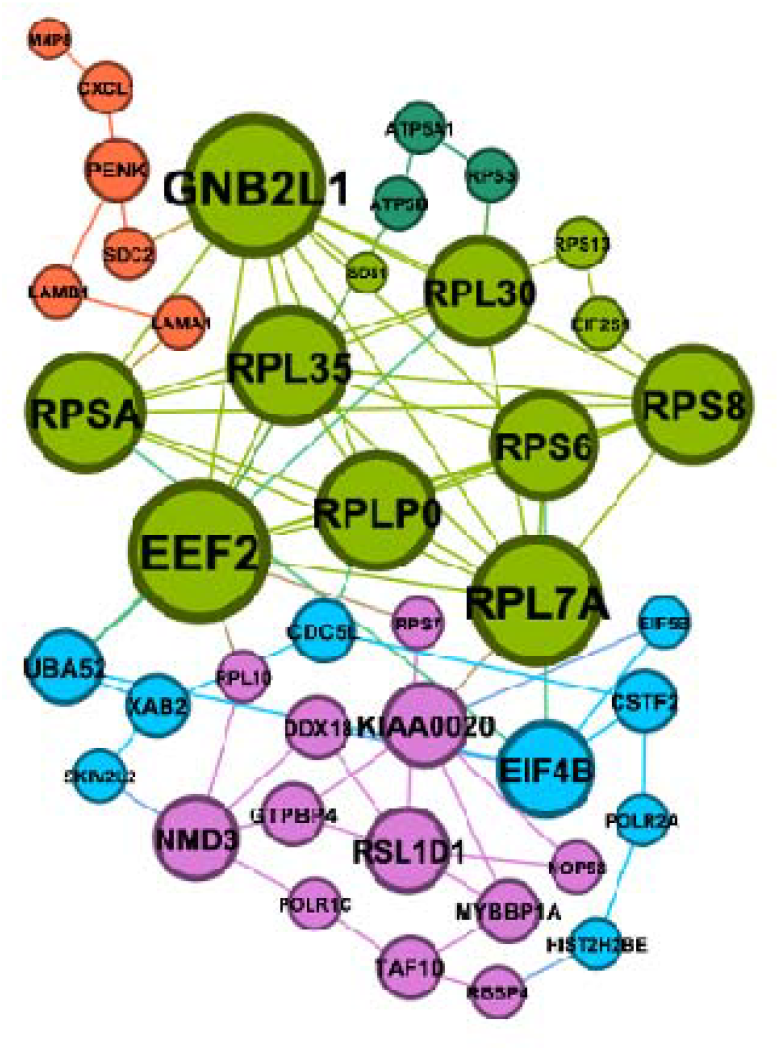
Network visualization of Cluster A, produced with Gephi. Different colors of nodes indicate modularity, a measure of the strength of the network graph’s division into clusters. The relative size of each node indicates its number of connections. Cluster A exhibits a module composed of several ribosome biogenesis hub genes.

**Figure 6.**
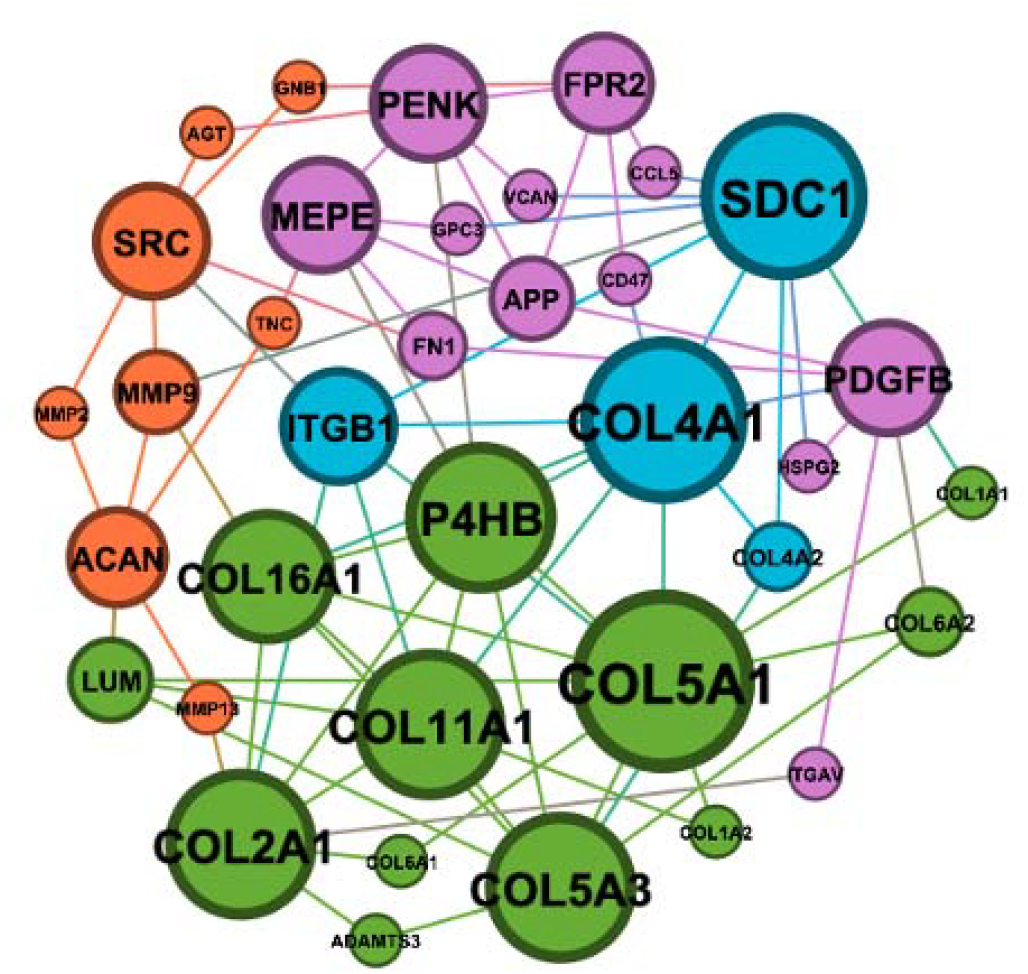
Network visualization of Cluster B, produced with Gephi. Cluster B exhibits a module composed of several collagen hub genes.

**Figure 7.**
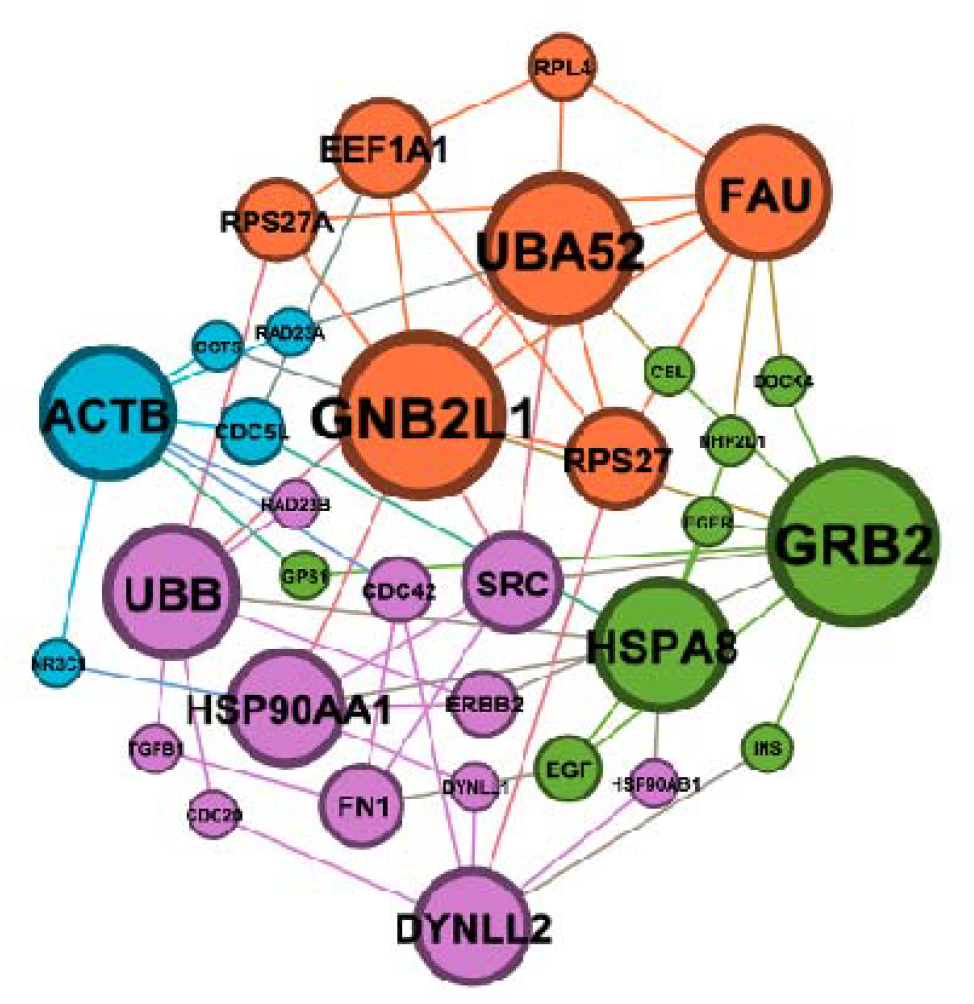
Network visualization of Cluster C, produced with Gephi. Cluster C exhibits several hub genes, including *GNB2L1, HSPA2, HSP90AA1, UBA52, UBB*, and *ACTB*.

**Figure 8.**
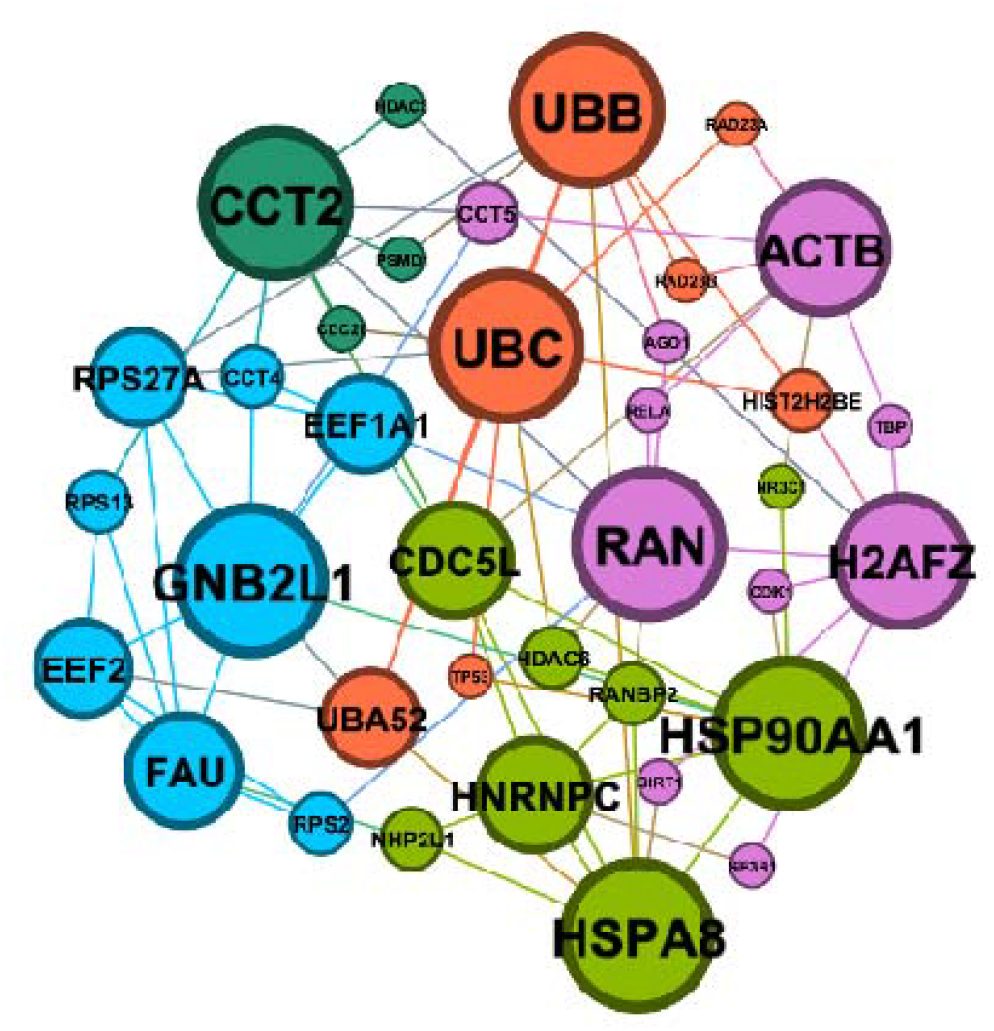
Network visualization of Cluster D, produced with Gephi. Cluster D exhibits several hub genes that are also present in Cluster C, including *GNB2L1, GRB2, HSPA2, HSP90AA1, UBA52, UBB*, and *ACTB*.

### Annotation

We annotated the four clusters with g:Profiler’s functional annotation tool, focusing on significant molecular functions, biological processes, cellular components, and KEGG pathways. Table 2 provides a summary of each cluster’s top 5 processes and pathways. Cluster A’s annotations were characterized by ribosome associated pathways and processes, as well as general protein synthesis regulation. These annotations reflect the network of ribosome biogenesis genes at the heart of the Cluster A network. Cluster B’s annotations were largely associated with collagen pathways, extracellular matrix organization, and cellular structure. Such annotations are highly connected to the network of collagen genes that formed a hub in the center of the Cluster B network. Cluster C and D produced similar annotations, characterized by more general pathways and processes involving protein binding regulation of protein metabolic activity.

**Table 2.**
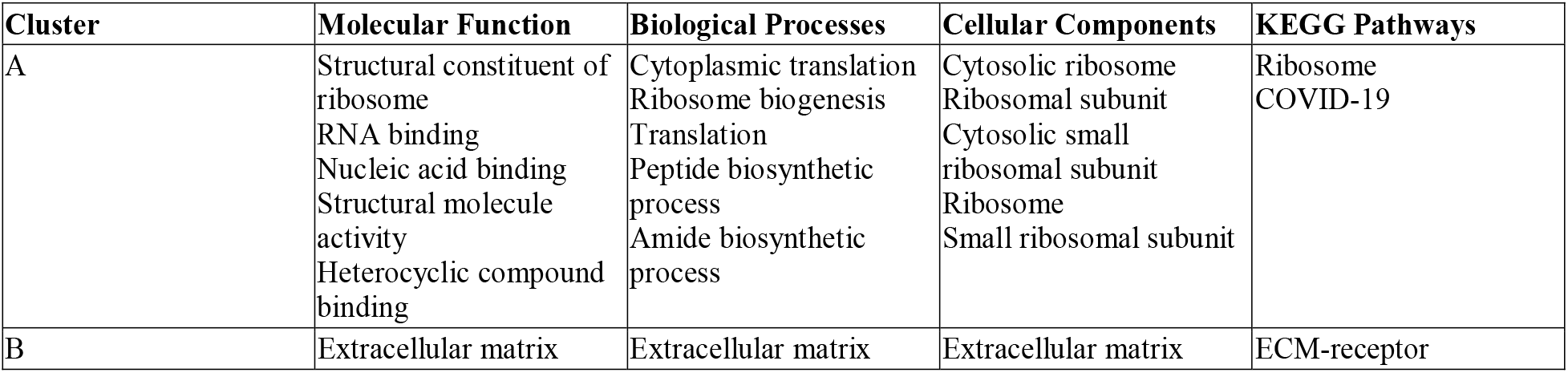
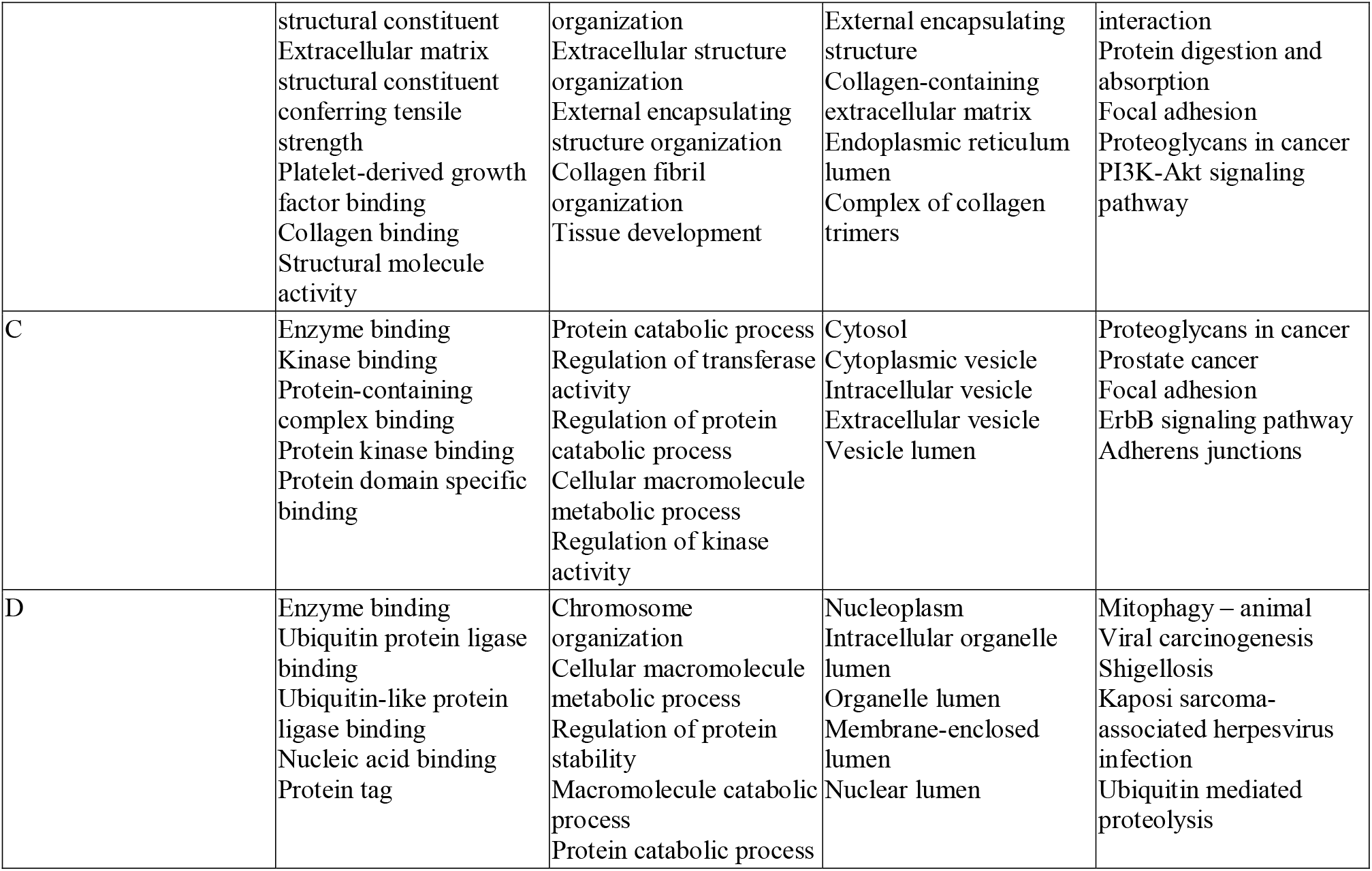
Summary of each cluster’s top 5 associated molecular functions, biological processes, cellular components, and KEGG pathways, as annotated by g:Profiler’s functional annotation tool.

### Separation Tests

A positive separation test score indicates topological separation of the disease modules, while a negative score indicates topological overlap between the disease modules. We observed a mixture of overlapping and separated networks. Clusters C and D demonstrated the most significant overlap with a separation score of -0.4112, indicating a high degree of topological overlap between the two networks in the STRING interactome. Clusters B and D had the most positive separation score of 0.7903, indicating a high degree of topological separation between the two networks. Table 3 summarizes the results of all 10 separation tests that were run.

**Table 3.** Summary of all 10 separation tests conducted. When a cluster is run against itself, the score provides a relative comparison for what a complete overlap for that network would look like.

|  | Cluster A | Cluster B | Cluster C | Cluster D |
| --- | --- | --- | --- | --- |
| Cluster A | -1.358974359 |  |  |  |
| Cluster B | 0.5267980743 | -1.176470588 |  |  |
| Cluster C | -0.07257473709 | 0.3087870384 | -1.129032258 |  |
| Cluster D | -0.09995049995 | 0.7092802338 | -0.4112693758 | -1.057142857 |

### Survival Analysis

Using outcome data on survival included with the TARGET dataset, we were able to analyze survival data for Clusters A and B. For Cluster A, 4 of the 6 patients died from disease, with a median of 30 months of survival. For Cluster B, 7 of the 13 patients died from disease, with a median of 54 months of survival. Of the remaining 194 high-risk patients from the TARGET dataset who were not in Clusters A or B, 126 died from disease, with a median of 42 months of survival. Calculated differences in survival were not significant, but the small sample sizes of Clusters A and B likely impacted the significance level. A Kaplan-Meier plot with Cluster A, Cluster B, and all other high-risk patients is depicted in Figure 9.

**Figure 9.**
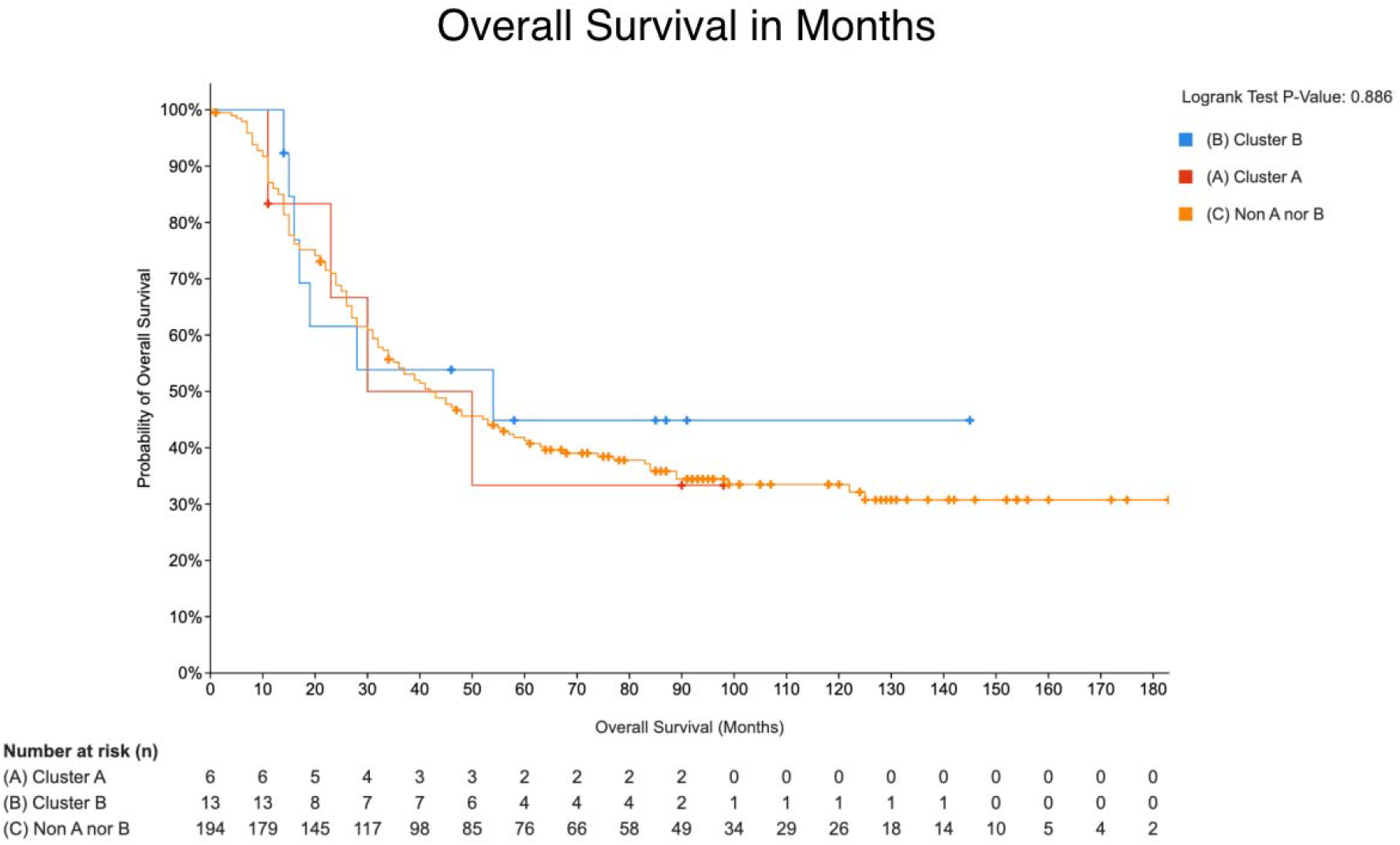
Kaplan-Meier plot of Cluster A, Cluster B, and remaining high-risk patients from the TARGET dataset by months. This KM plot was generated using cBioPortal.

The GSE49711 dataset solely provided binary data on death from disease. For Cluster C, 33 of the 65 patients died from disease (50.8%). Of the remaining 110 high-risk patients not included in Cluster C, 57 died from disease (51.8%).

## Discussion

The shared PPI networks identified in each cluster present potential targets for novel therapies in the treatment of the specific subgroup of high-risk *MYCN* amplified neuroblastoma patients.

### Cluster A: Targeting Ribosome Biogenesis Networks

In Cluster A, a network of highly connected ribosome biogenesis genes, including *RPL7A, RPL30, RPL35*, and *RPS8*, was identified as specific to *MYCN* amplified patients. This finding aligns with existing literature that highlights *MYCN* as a regulator of ribosome biogenesis and protein synthesis (15, 16). Notably, prior to filtering out ribosome and histone-associated genes in the GSE49711 and GSE89413 datasets, an overcrowding of ribosome biogenesis genes was observed. The overexpression of these genes suggests their potential as therapeutic targets. A study by Hald et al. demonstrated that ribosome biogenesis inhibitors can repress the growth of *MYCN* amplified neuroblastoma, indicating the therapeutic potential of targeting this network (17).

### Cluster B: Targeting Collagen Catabolism and Matrix Metallopeptidase Networks

Cluster B revealed three fibrillar and two non-fibrillar collagen genes—*COL11A1, COL16A1, COL2A1, COL5A3*, and *COL6A2*—unique to high-risk *MYCN* amplified patients. These genes are associated with extracellular matrix (ECM) organization, a factor known to influence *MYCN* expression via ECM stiffness (18). Additionally, these genes are linked to increased activation of the PI3K-AKT pathway, which is correlated with poor prognosis in neuroblastoma (19). Given that collagen-activated PI3K-AKT signaling enhances tumor growth in other cancers, these findings suggest a similar metastatic mechanism may be at play in *MYCN* amplified neuroblastoma (20, 21). The matrix metallopeptidase gene network identified in Cluster B is also noteworthy. *MMP2* and *MMP13* are unique to *MYCN* amplified patients, while *MMP9* is inferred from the interactome. Previous studies have associated overexpression of *MMP2* with poor outcomes in neuroblastoma (22). These results suggest that further research should explore whether these matrix metallopeptidase genes and their role in extracellular matrix remodeling represent a unique pathway for tumor metastasis in *MYCN* amplified neuroblastoma.

### Cluster C: Targeting Heat Shock Protein and Hub Gene Networks

Cluster C is characterized by an upregulated heat shock protein network, including *HSPA8, HSP90AA1*, and *HSP90AB1*, among *MYCN* amplified patients. This supports existing studies that propose heat shock protein inhibitors as effective anticancer agents (23). Additionally, Cluster C features several hub genes—*EEF1A1, UBB, GNB2L1*, and *ACTB*—whose roles in cancer have been highlighted in other studies. Downregulation of *EEF1A1* has been suggested as a breast cancer therapeutic target, whereas upregulation of *UBB* and upregulation of *GNB2L1* have separately been proposed as potential gastric cancer therapeutic targets (24-26). Upregulation of *ACTB* has been implicated as a biomarker for poor prognosis and immune infiltration in a pan-cancer study (27). The highly interconnected nature of these genes within the Cluster C network warrants further investigation into their potential synergistic roles in promoting metastasis in *MYCN* amplified neuroblastoma.

### Cluster D: Validation with Cell-Line Data

Cluster D, which is composed of tumor-derived cell-lines, exhibited gene networks strikingly similar to those in Cluster C. The strong connection between *HSPA8* and *HSP90AA1* among *MYCN* amplified cell lines reinforces the potential of heat shock protein inhibitors as a therapeutic strategy for *MYCN* amplified neuroblastoma. Importantly, the cell lines in Cluster D offer a valuable resource for in-vitro experimentation to test novel targeted therapies.

#### Survival Analysis: Implications, Limitations, and Future Directions

The survival analysis conducted in this study provides important insights into the clinical significance of the identified PPI networks in high-risk *MYCN* amplified neuroblastoma patients. While the clusters do not predict groups of patients with worse overall survival, they do identify distinct populations within high-risk patients. However, it is essential to acknowledge certain limitations, particularly concerning the sample size in Clusters A and B, which may influence the interpretation of these findings.

In Cluster A, the network of ribosome biogenesis genes is associated with a median survival of 30 months, indicating a potential therapeutic target. However, this cluster is composed of only 6 patients, with just 4 patients who did not survive the disease. The small sample size limits the generalizability of these findings, and further studies with larger cohorts are necessary to validate the role of ribosome biogenesis genes as reliable biomarkers and therapeutic targets. Despite this limitation, the consistency of these findings with existing literature suggests that targeting this pathway could still be a promising area for future research and clinical trials aimed at improving survival outcomes in this subgroup.

Similarly, Cluster B contains only 13 patients with a median survival of 54 months. While the observed associations between the collagen catabolism and matrix metallopeptidase networks are intriguing and align with known mechanisms of extracellular matrix organization and PI3K-AKT signaling in tumor progression, the limited sample size necessitates caution in drawing definitive conclusions. Larger studies are needed to confirm these results and to better understand the potential for targeting these pathways in improving survival for *MYCN* amplified neuroblastoma patients.

The findings in Clusters C and D, the latter being based on cell-line data, highlight the significance of heat shock protein networks and associated hub genes as therapeutic targets. The substantial overlap between these clusters, coupled with the high mortality rate observed, underscores the potential significance of developing targeted therapies for these networks. However, as with Clusters A and B, larger and more diverse patient cohorts will be essential to validate these findings and translate them into clinical practice.

In conclusion, while our data suggest promising directions for therapeutic intervention, the small sample sizes in Clusters A and B represent a significant limitation. Future research should aim to include larger patient cohorts to confirm the prognostic value of these PPI networks. Moreover, integrating these findings with additional clinical variables and conducting longitudinal studies will help refine their relevance in clinical settings. By addressing these limitations, there is strong potential to enhance survival rates and improve the management of high-risk *MYCN* amplified neuroblastoma.

